# Visual evidence for the recruitment of four enzymes with RNase activity to the *Bacillus subtilis* replication forks

**DOI:** 10.1101/2022.09.13.507877

**Authors:** Rebecca Hinrichs, Peter L. Graumann

## Abstract

Removal of RNA/DNA hybrids for the maturation of Okazaki fragments on the lagging strand, or due to misincorporation of ribonucleotides by DNA polymerases, is essential for all types of cells. In prokaryotic cells such as *Escherichia coli*, DNA polymerase 1 and RNase HI are supposed to remove RNA from Okazaki fragments, but many bacteria lack HI-type RNases, such as *Bacillus subtilis*. Here, four proteins have been shown to be able to remove RNA from RNA/DNA hybrids *in vitro*, but their actual contribution to DNA replication is unclear. We have studied the dynamics of DNA polymerase A (similar to Pol 1), 5’->3’ exonuclease ExoR, and the two endoribonucleases RNase HII and HIII in *B. subtilis* using single molecule tracking. We found that all four enzymes show a localization pattern similar to that of replicative DNA helicase. By scoring the distance of tracks to replication forks, we found that all four enzymes are enriched at DNA replication centers. After inducing UV damage, RNase HIII was even more strongly recruited to the replication forks, and PolA showed a more static behavior, indicative of longer binding events, whereas RNase HII and ExoR showed no response. Inhibition of replication by HPUra clearly demonstrated that both RNase HII and RNase HIII are directly involved in replication, with RNase HIII playing a major role. We found that the absence of ExoR increases the likelihood of RNase HIII at the forks, indicating that substrate availability rather than direct protein interactions may be a major driver for the recruitment of RNases to the lagging strands. Thus, *B. subtilis* replication forks appear to be an intermediate between *E. coli* type and eukaryotic replication forks and employ a multitude of RNases, rather than any dedicated enzyme for RNA/DNA hybrid removal.

## Introduction

RNA maturation, RNA degradation, and RNA turnover are essential processes for all kinds of cells. Enzymes known as ribonucleases (RNases) are crucial for these processes (Williams & Kunkel, 2014), and are usually present in many different versions per cell. During DNA replication, RNases play a major role in the maturation of Okazaki fragments on the discontinuous, lagging strand, by removing the arising DNA/RNA hybrids made by DNA primase (Li & Breaker, 1999). In addition, even though DNA polymerases have a much higher affinity for desoxy ribonucleotides (dNTPs) than ribonucleotides, the RTP pools are in large excess of those of dNTPs, wherefore ribonucleo-monophosphates (rNMPs) are incorporated about every 2.3 kb in *E. coli* cells (Yao *et al*., 2013). Loss of removal of for rNMPs leads to an increase in mutation frequency.

A general differentiation can be made between endoribonucleases and exoribonucleases. RNase E or RNase Y are major endoribonucleases that initiate RNA turnover, which is then taken over by exonucleases (Bechhofer & Deutscher, 2019). The latter process RNA molecules that result in 3’ or 5’ terminal release of nucleotide residues. A 3’ to 5’ exoribonuclease in *Bacillus subtilis* is exemplified by polynucleotide phosphorylase (PNPase), while magnesium-dependent ExoR or RNase J1 are 5’ to 3’ exonucleases (Li & Deutscher, 2004, Bechhofer & Deutscher, 2019). In addition to the classical ribonucleases, several other enzymes also exhibit exonucleolytic functions on RNA/DNA hybrids, like the DNA polymerase Pol I or PolA, which exhibit 5’->3’ exonuclease activity in *E. coli* or *B. subtilis*, respectively (Duigou *et al*., 2005). Pol I and PolA are thought to contribute to RNA removal at replication forks in bacteria, in conjunction with or in addition to different RNases. During DNA replication in eukaryotes, two RNases were discovered, which belong to the RNase H family: RNase H1 and RNase H2 hydrolyze RNA from RNA/DNA hybrids (Cerritelli & Crouch, 2009). Prokaryotes possess one or two of three H-type RNases, RNase HI, HII, and RNase HIII. In *E. coli*, RNase HI hydrolyzes RNA-DNA hybrids, which contain polymers of four or more ribonucleotides (Ohtani *et al*., 1999b). RNase HII is characterized by hydrolyzing at the 5′ to a single ribonucleotide, i.e. it acts as an endonuclease, unlike RNase HI. Also, it hydrolyzes at the 5′ to the ribonucleoside monophosphate (Yao *et al*., 2013) Thus, it is important to initiate removal of incorporated rNMPs from the genome, to ensure genomic integrity (Clark & Kunkel, 2010). RNase HIII has a high enzymatic similarity to RNase HI (Ohtani *et al*., 1999a). Bacteria are considered to generally have two RNase H enzymes, which in *Bacillus subtilis* are RNase HII and RNase HIII (Kochiwa *et al*., 2007). Most endoribonucleases cleave RNA in the presence of divalent cations, resulting in fragments containing 3′-hydroxyl and 5′-phosphate termini (Li & Deutscher, 2004), including RNase HII and RNase HIII.

During DNA replication in *B. subtilis*, continuous synthesis of the leading strand occurs by polymerase PolC. For lagging strand synthesis, primase DnaG synthesizes RNA primers, which are extended by DNA polymerase DnaE (Sanders *et al*., 2010, Dervyn *et al*., 2001). DnaE is thought to only extend by few bases, and then hand over to PolC (Seco & Ayora, 2017), which finishes the 1-2 kb Okazaki fragments (Ogawa & Okazaki, 1980). Indeed, 2000-4000 fragments per 4.6 Mb chromosome could be identified in *E. coli* for whole chromosome replication (Su’etsugu & Errington, 2011). Removal of RNA primers is important for DNA stability (Williams & Kunkel, 2014), and lagging strand synthesis is terminated by ligase LigA sealing remaining single strand gaps. As pointed out above, DNA polymerase PolA shows 5’-3’ exonuclease activity and could be detected together with RNase HIII and ExoR in *in vitro* studies for the maturation of Okazaki fragments in *B. subtilis* (Randall *et al*., 2019). Also, as stated above, RNase HII appears to be responsible for the removal of individual rNMPs that are incorporated into DNA by DNA polymerase during DNA replication (Yao *et al*., 2013, Schroeder *et al*., 2017). Similarly, PolA, as well as ExoR could be determined to be involved in the repair of DNA damage due to UV (Hernández-Tamayo *et al*., 2019). Interestingly, temperature-sensitive phenotypes were discovered in different deletions combinations. It was shown that the deletion of *rnhC* and *polA*, as well as the deletion of *rnhC* and *exoR*, lead to lethality under 25°C growth conditions (Randall *et al*., 2019), while a double mutation (*rnhB, rnhC*) leads to poor growth (Yao *et al*., 2013). These findings point to a possible role of these proteins at *B. subtilis* replication forks, but other explanations for synthetic phenotypes are possible. In order to obtain definite proof for enzymatic activity of the four mentioned proteins at replication forks, or at sites all over the chromosome distinct from replication events, we monitored single molecule dynamics, to test if replication forks represent sites of frequent stops for different enzymes having RNase activity.

## Materials and Methods

### Strain construction

For the construction of the mV /mNeo fusions in *B. subtilis*, the integration plasmid pSG1164 was used. Using pSG1164, the corresponding fluorophore is integrated through a single crossover c-terminal to the original locus (Lucena *et al*., 2018). For integration, a 500 bp homolog of the C-terminus of the gene of interest must be cloned using Gibson Assembly (Gibson *et al*., 2009) into the vector next to the linker and mV/ mNeo sequence. Used primers have a homologue 25 bp overhang and are listed in Table 1. The transformed plasmids were extracted using a kit (New England BioLabs). The deletion strains are based on *B. subtilis* BG214 and containing RNase HIII-mV or ExoR-mV constructs and were generated by transformation of cells with chromosomal DNA from *B. subtilis* 168 *ΔrnhC::kan trpC2, ΔexoR::kan trpC2*, obtained from the *Bacillus* Genetic Stock Center (Columbus, Ohio) (Koo *et al*., 2017). The chromosomal DNA was extracted using a kit (innuPREP Bacteria DNA Kit, Analytik-Jena).

**Table 1:** Strains or plasmids used in this study

| Strain or Plasmids | Relevant features | Reference or source |
| --- | --- | --- |
| *Bacillus subtilis* |  |  |
| BG214 | Wild type | gift from Juan C. Alonso |
| PG3977 | BKK40880 (ΔexoA::kan trpC2) | (Koo *et al.* 2017) |
| PG3777 | BKK28620 (ΔrnhC::kan trpC2) | (Koo *et al.* 2017) |
| PG4345 | *rnhB*-mV<sup>cmR</sup> | This study |
| PG4346 | *rnhB*-mNeo<sup>cmR</sup> *dnaX*-CFP<sup>specR</sup> | This study |
| PG4347 | *rnhC*-mV<sup>cmR</sup> | This study |
| PG4348 | *rnhC*-mNeo<sup>cmR</sup> *dnaX*-CFP<sup>specR</sup> | This study |
| PG4349 | Δ*exoR*::kan trpC2 *rnhC*-mV<sup>cmR</sup> | This study |
| PG4350 | *exoR*-mV<sup>cmR</sup> | This study |
| PG4351 | *exoR*-mNeo<sup>cmR</sup> *dnaX*-CFP<sup>specR</sup> | This study |
| PG4352 | Δ*rnhC*::kan trpC2 *exoR*-mV<sup>cmR</sup> | This study |
| PG4353 | *polA*-mNeo<sup>cmR</sup> *dnaX*-CFP<sup>specR</sup> | This study |
| PG4374 | *polC*-mNeo<sup>cmR</sup> *dnaX*-CFP<sup>specR</sup> | This study |
| PG3173 | *dnaX*-CFP<sup>specR</sup> | (Lindow *et al.* 2002) |
| PG3730 | DH5α pSG1164-mVenus, expression Vektor, Amp<sup>R</sup>, Cm<sup>R</sup> | (Lucena *et al.* 2018) |
| 131 | pSG1164 *rnhB*-mVenus, integration Vector, Amp<sup>R</sup>, Cm<sup>R</sup> | This study |
| 132 | pSG1164 *rnhB*-mNeonGreen, integration Vector, Amp<sup>R</sup>, Cm<sup>R</sup> | This study |
| 133 | pSG1164 *rnhC*-mVenus, integration Vector, Amp<sup>R</sup>, Cm<sup>R</sup> | This study |
| 134 | pSG1164 *rnhC*-mNeonGreen, integration Vector, Amp<sup>R</sup>, Cm<sup>R</sup> | This study |
| 135 | pSG1164 *polA*-mNeonGreen, integration Vector, Amp<sup>R</sup>, Cm<sup>R</sup> | This study |
| 136 | pSG1164 *exoR*-mNeonGreen, integration Vector, Amp<sup>R</sup>, Cm<sup>R</sup> | This study |
| 137 | pSG1164 *exoR*-mVenus, integration Vector, Amp<sup>R</sup>, Cm<sup>R</sup> | This study |

**Table 2:**
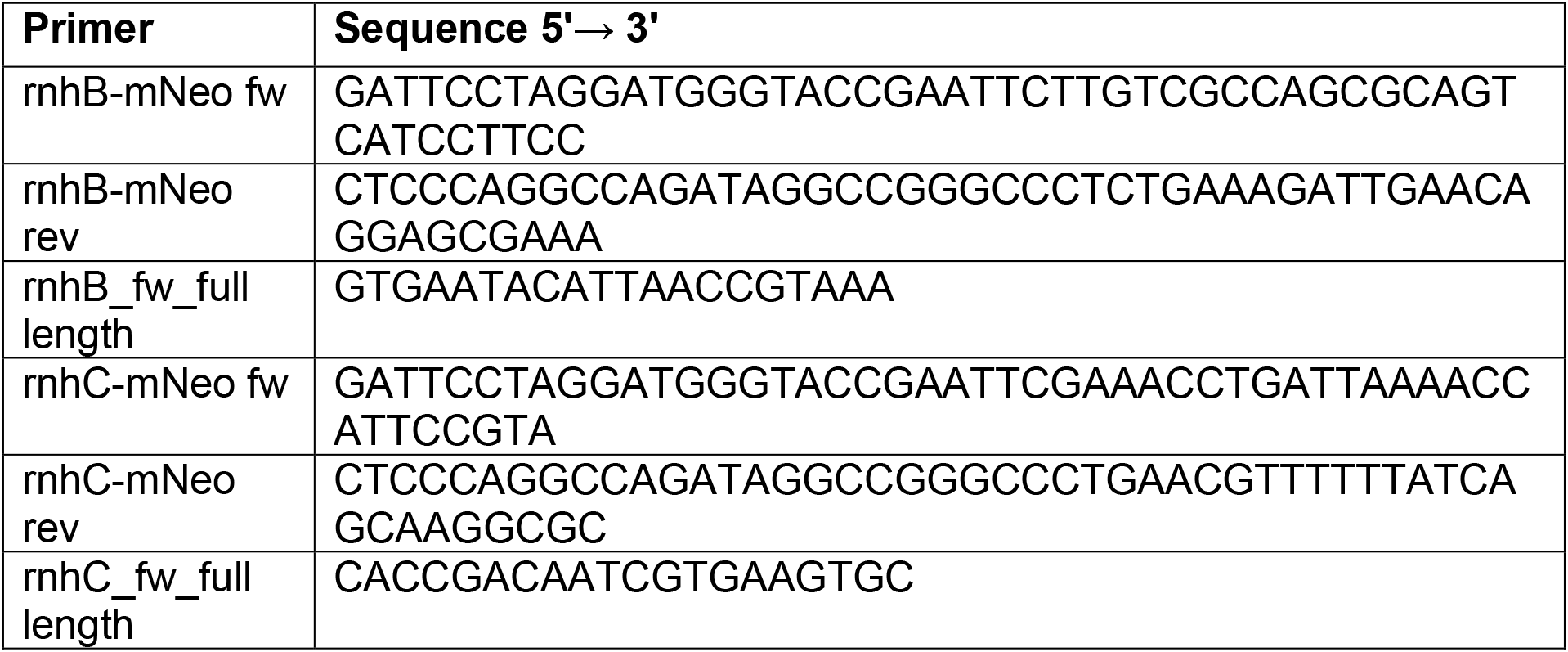
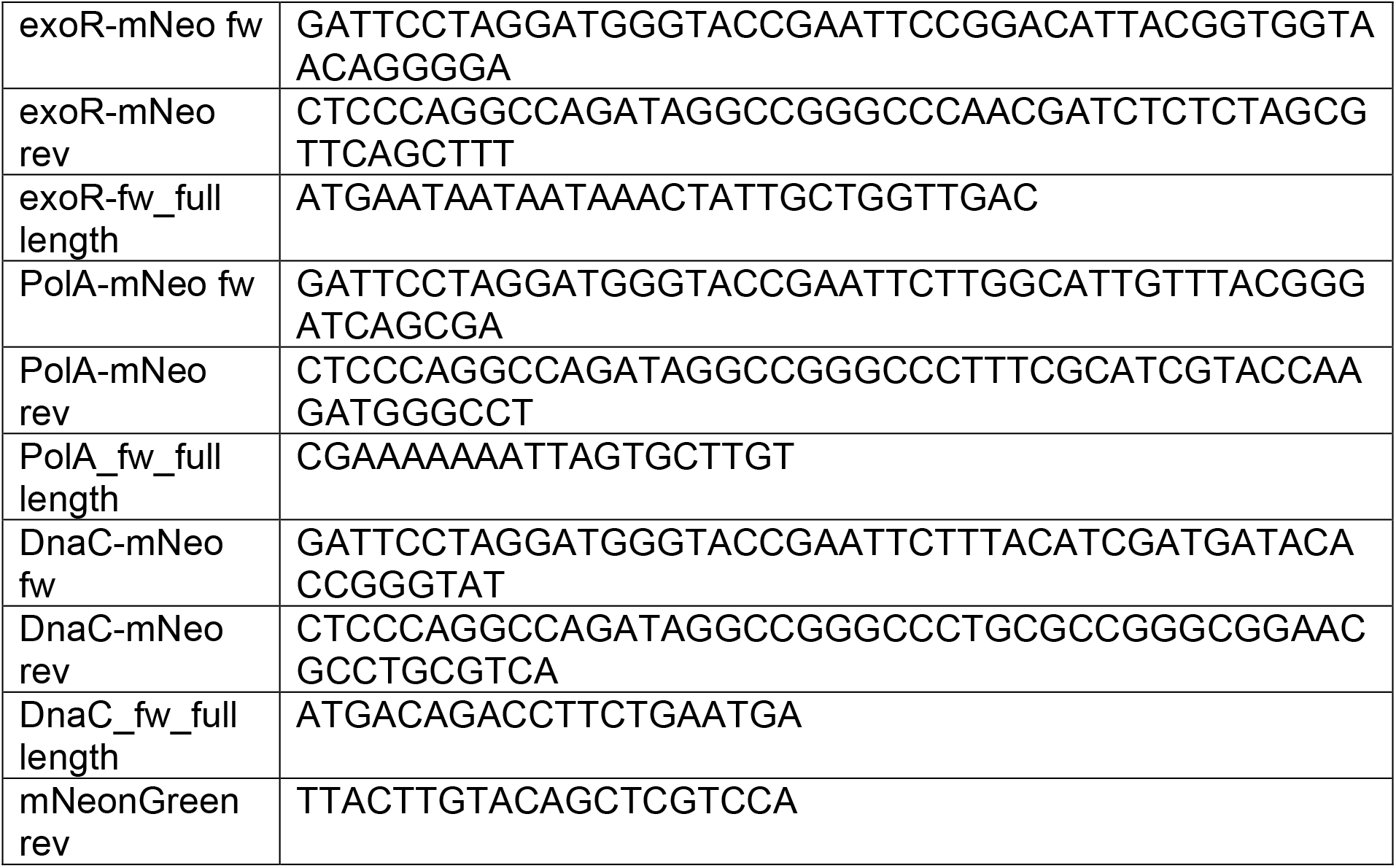
List of Oligonucleotide.

### Growth conditions

All plasmids, strains and oligonucleotides used are listed in tables in the Supplement. *E. coli* strains were cultured in LB (lysogenic broth) medium at 37°C. Likewise, LB medium was used for microscopy (SMT). For this experiment, the *B. subtilis* strains were cultured in LB at 30°C. When needed, antibiotics were added at the following concentrations: ampicillin 100 μg/ml, chloramphenicol 5 μg/ml, kanamycin 30 μg/ml, 6(p-hydroxyphenylazo)-uracil (HPUra) 15 mg/ml. 0.5% of xylose was added from a 50% sterile filtrated stock solution in ddH2O.

### Western blot

The samples were harvested from the exponential growth phase and digested by lysozyme (proceed according to (Hinrichs *et al*., 2022)). The detection was performed with a primary polyclonal α GFP-tag antibody (1:5000)/ mNeonGreen-tag (1:4000), and secondary antibody goat-anti-Rabbit-IgG, peroxidase-conjugated (1:100,000) (Sigma-Aldrich).

### Fluorescence microscopy

Microscopy was performed in LB medium with a prior cultivation of 30°C, 200 rpm. Cells were analized in the exponential growth phase. For wide-field epifluorescence microscopy, a Zeiss Observer A1 microscope (Carl Zeiss) with an oil immersion objective (100 x magnification, 1.45 numerical aperture, alpha PlanFLUAR; Carl Zeiss) was used. The images were recorded with a charge-coupled-device (CCD) camera (CoolSNAP EZ; Photometrics) and an HXP 120 metal halide fluorescence illumination with intensity control (Hinrichs *et al*., 2022). For the sample preparation a round coverslips (25 mm, Marienfeld) was used and covering 5 μl cell culture with a 1.5 % agarose pad. The agarose pads were made with water by sandwiching 100 μl of the melted agarose between two smaller coverslips (12 mm, Menzel). Images were processed using ImageJ (Schindelin *et al*., 2012).

### Single-molecule tracking (SMT)

Individual molecules were tracked using custom-made slim-field setup on an inverted fluorescence microscope (Nikon Eclipse Ti-E, Nikon Instruments Inc.). An EMCCD camera (ImagEM X2 EM-CCD, Hamamatsu Photonics KK) was used to ensure high-resolution detection of the emission signal, resulting in a calculated resolution of the position of the molecule down to 20 nm. The central part of a 514 nm laser diode (max power 100 mW, TOPTICA Beam Smart) was used with up to 20% of the intensity (about 160 W cm-2 in the image plane) to excite samples, fused to mNeonGreen (using laser filter set BrightLine 500/24, dichroic mirror 520 and BrightLine 542/27), by focusing the beam onto the back focal plane of the objective. A CFI Apochromat objective (TIRF 100 x Oil, NA 1.49) was used in the setup (Oviedo-Bocanegra *et al*., 2021). For the analysis, a video of 3000 frames at 20 ms was recorded, of which 1000 starting after 200 to 300 frames, dependent on the time point when single molecule levels was reached due to bleaching, were used for the analysis. Software Oufti (Paintdakhi *et al*., 2016) was used to set the necessary cell meshes. Utrack (Jaqaman *et al*., 2008) was employed for automatic determination of molecule trajectories. Data analysis was carried out using software SMTracker 2.0 (Oviedo-Bocanegra *et al*., 2021, Dersch *et al*., 2020).

## Results

### PolA, ExoR and RNase HII show nucleoid localization

To investigate the localization of PolA, ExoR, RNase HII, and RNase HIII *in vivo*, C-terminal mVenus or mNeonGreen fusions were created and integrated at the original locus. The localization of the replication forks was visualized using a DnaX-CFP (DNA polymerase III, part of the clamp-loader complex) allele in the same strain. To ensure that the fused proteins are expressed in full length, a Western blot was made against the corresponding fluorophore (Fig. S1/S2). The experiments showed full-length expression in all cases.

Using epifluorescence microscopy (Fig. 1), we found that ExoR-mNeo, PolA-mNeo and RNase HII-mNeo show a clear staining pattern comparable to a DAPI stain of the nucleoids in the cell (Pediaditakis *et al*., 2012). The fusion of RNase HIII showed a diffuse localization throughout the cells (Fig. 1). The CFP channel provides information about the localization of the replication forks: the DnaX-CFP fusion showed distinct foci in the cell (1-2 per cell). The merge of both channels shows the colocalization of the proteins relative to DnaX, showing clear overlap of signals.

**Figure 1:**
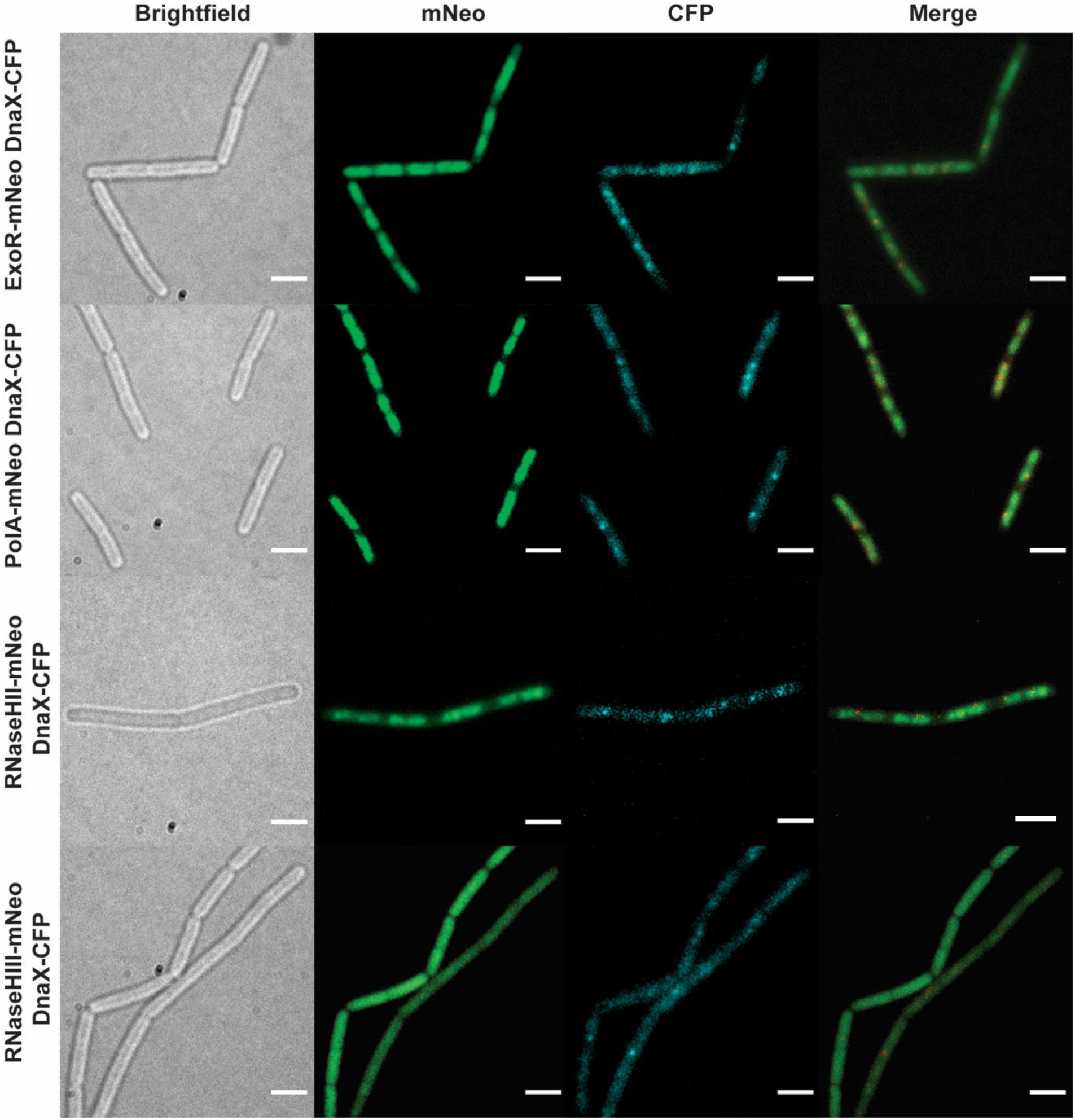
Epifluorescence microscopy of fusion strains. The image shows the brightfield image, the mNeonGreen of the fusions, DnaX-CFP for replication fork localization, and a corresponding merge. PolA-mNeo, ExoR-mNeo, RNase HII-mNeo and, RNase HIII-mNeo were colocalized with DnaX-CFP. Scale bar 2 μm.

### Single molecule tracking reveals distinctive patterns of motion for RNases

Epifluorescence experiments could not reveal an association of any RNase or of PolA with replication forks. To obtain a better understanding of protein dynamics and increased spatiotemporal resolution, we used single-molecule tracking (SMT) (Oviedo-Bocanegra *et al*., 2021). The method employs a beam of a 514-nm laser diode is expanded by a factor of 20, and the central part is focused on the rear focal plane of the 100 x A = 1.49 objective. SMT allows visualization of events of molecules located at a defined subcellular location with an accuracy of 40 nm or less (Dersch *et al*., 2020). SMT was performed with 20 ms stream acquisition. To determine the area of the cell to be detected, cell-meshes are set by using Oufti (Paintdakhi *et al*., 2016) and trajectories were determined by u-track (Jaqaman *et al*., 2008). Tracks of only 5 steps and more were used. Analysis of the resulting data, all from biological triplicates, was done with the SMTracker 2.0 (Oviedo-Bocanegra *et al*., 2021).

Figure 2A shows the heat maps of the given proteins to visualize the localization of molecule tracks in the cell. For this, we projected all tracks from the biological replicates into a cell with an average size of 3 × 1 μm. The distribution of tracks is indicated by a color shift from yellow (low probability) to black (highest probability). The intensity of the maps created is adapted to each other. To be able to compare localization patterns with already known replication proteins, a fusion of DnaC (DNA helicase) was included (Bin *et al*., 2013). Similar to the epifluorescence images (Fig 1), the localization of PolA, ExoR, and RNase HII fusions are clearly seen on the nucleotides, with a concentration to the central parts of the nucleoids, very similar to DnaC. The localization of RNase HIII is very similar, but with a lower intensity gradient from central parts to sites surrounding the nucleoids. RNase HIII most strongly resembled the heat map of DnaC (Fig 2A). Contrarily, RNase J2, component of the RNA degradosome, which is mostly membrane-associated, showed strong accumulation towards the cell periphery and was depleted from many central positions. These data support the notion that RNase HII and HIII are associated with removal of RNA/DNA hybrids from the DNA, and may be associated with replication.

**Figure 2:**
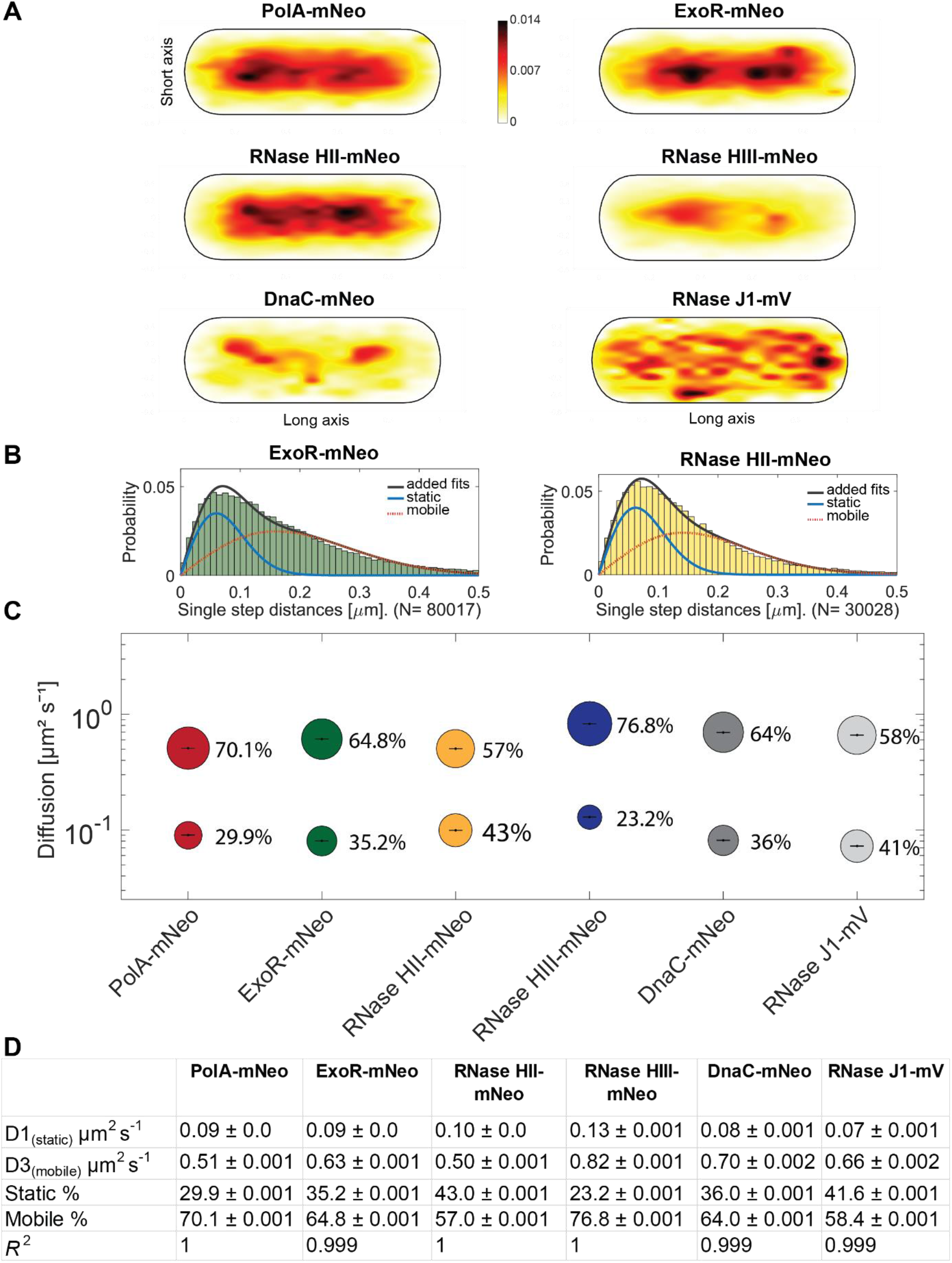
Single molecule analyses of PolA-mNeo, ExoR-mNeo, RNase HII-mNeo, RNase HIII-mNeo and, DnaC-mNeo. (A) Heat maps of single-molecule localization of replication proteins in a medium-size *Bacillus subtilis* cell. The distribution of tracks is indicated by a color shift from yellow (low probability) to black (highest probability). (B) Jump distance analyzes of ExoR-mNeo and RNase HII-mNeo. The two rayleigh fits display a two-population fit (static in blue, mobile in red and, added fits in black). (C) Bubble blots show diffusion constants of replication Proteins and fractions sizes for static and mobile molecules. (D) Diffusion constants and percentages of static and mobile molecule fractions. Values were fitted using non-linear least-square fitting, R2 values for each condition are stated.

We next determined diffusion constants of the proteins using squared displacement analyses (SQD), shown in Figures 2B and 2C. SQD analysis can be used to determine the average diffusion constant of a molecule, as well as to analyse if a single or several populations with different diffusion constants exist; if several, the size of the populations can be determined. For PolA, ExoR and RNases, a two-population Rayleigh fit was used, which explained the observed distribution of tracks very well. This analysis suggests the existence of a slow mobile/static and a high mobile population, most likely freely diffusive molecules. The bubble blot (Fig. 2C) visualizes diffusion constants and sizes of the two assumed populations, revealing that all proteins have slow-mobile fractions whose diffusion constants are in a similar range, one that has been described for tight DNA-binding events. In case of DnaC, this is tight hexamer formation ahead of replication forks. ExoR, RNase HII and, DnaC have the largest slow mobile/static fractions, RNase HIII has the largest diffusive population (Fig 2B). Thus, a quarter to a third of molecules appears to be engaged in a substrate-bound form. For detailed numbers, please also consult table 1. Although informative, these data still do not provide an answer if RNase activity of the studied proteins is associated with replication forks *in vivo*, other than for PolA and ExoR, where spatial connection to replication has been shown before (Hernández-Tamayo *et al*., 2019).

### All four enzymes showing RNase activity feature close spatial proximity of motion to are frequent arrests at replication forks

A convincing argument that an enzyme takes place in a reaction at a defined subcellular space in proximity of motion at or close to that site. We used a tool in SMTracker 2.0 that allows to score proximity of molecule trajectories close to sites in the cell that can be defined e.g. by localizing a protein complex using a protein fusion having a different fluorescent colour. Fig. A shows an example of two cells in which the position of replication forks has been determined by acquiring an image in the CFP channel, detecting DnaX-CFP, a component of the clamp loading complex. Blue tracks reveal freely diffusive trajectories, red tracks trajectories staying within a radius of 106 nm for at least 5 time points, i.e. molecules showing confined diffusion, likely due to binding events. These are close to DnaX-CFP signals, and thus at replication forks.

Green trajectories reveal events of transition between free diffusion and confined motion, logically being close to events of confined motion. Thus, our analyses can capture events of transitions from free diffusion to DNA binding, and release from DNA binding. The lower right cell in Fig. 3A likely contains a second replication fork close to the two events of confined motion for RNase HII, which was not captured in the CFP channel. As a definition of what “close to replication forks” means, we tracked DnaC-mNeo relative to DnaX-CFP. Fig. 3B shows a roughly Gaussian distribution of tracks with a centre at about 400 nm. The relatively large deviation from “0” is partially due to a change in filters and illumination between taking the CFP image by epifluorescence and by tracking DnaC-mNeo molecules by SMT, because replication forks are quite mobile (Monahan *et al*., 2014). In addition, replication forks can only be located with a resolution of 250 nm, while SMT has a localization error of less than 40 nm. Fig. 3B contains a second peak at about 1.4 μm, likely DnaC-mNeo moving close to replication forks that have not been identified in epifluorescence.

**Figure 3:**
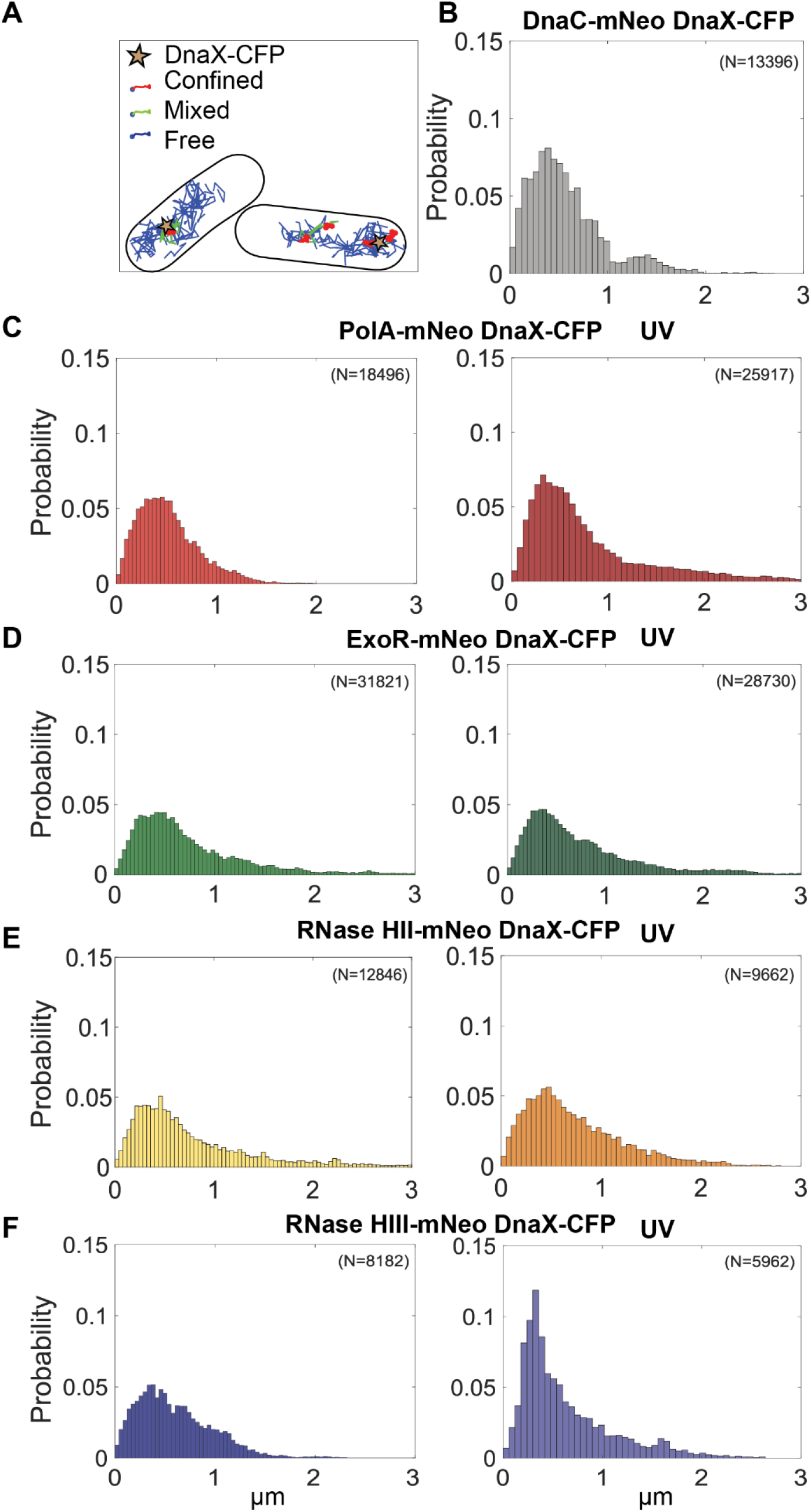
Distance measurement of potential replication proteins to DnaX-CFP (replication fork). Panel A shows an example for the confinement analyses of RNase HII-mNeo. The marker for the replication fork is DnaX-CFP, expressed from the original gene locus. At least 50 cells were measured. The number of tracks measured is indicated (N). The probability of detection close to DnaX-CFP foci is shown in relation to the distance in micrometers (μm). The above proteins are described in A-E.

For PolA-mNeo, the distribution is between 0 μm and 2 μm distance to the DnaX-CFP signal is centered at a distance of about 0.4 μm, and thus similar to DnaC-mNeo. The same is true for all three other enzymes investigated, indicating that all frequently arrest at replication forks in their search for DNA/RNA hybrids.

The above data revealed that PolA, ExoR, RNase HII and HIII show high probability of motion close to sites of DNA replication. These analyses do not quantify the extent of motion, or rather arrest, at replication forks. We searched for further evidence for the presence of four enzymes with RNase activity by confinement analysis in relation to the replication forks (Fig 4). For this, scored for confined tracks, i.e. tracks that stay within three times the localization precision of this work, for 5 or more time points. We analyzed only cells with distinct DnaX-CFP signals and counted how many cells had confined tracks immediately at a replication fork (Fig. 4A). As we used a tracking time of 1000 frames, confinement analyses shows percentage of cells having molecules arresting at DnaX-CFP foci within a 20 seconds time frame. Interestingly, all enzymes (PolA (71.4%), ExoR (72.7%), RNase HII (76%), and RNase HIII (72.1%) showed confined tracks at replication forks in more than 70% of cells (Fig. 4B and 4C). As reference, for DNA helicase reference, a similar percentage of more than 78% could be determined.

**Figure 4.**
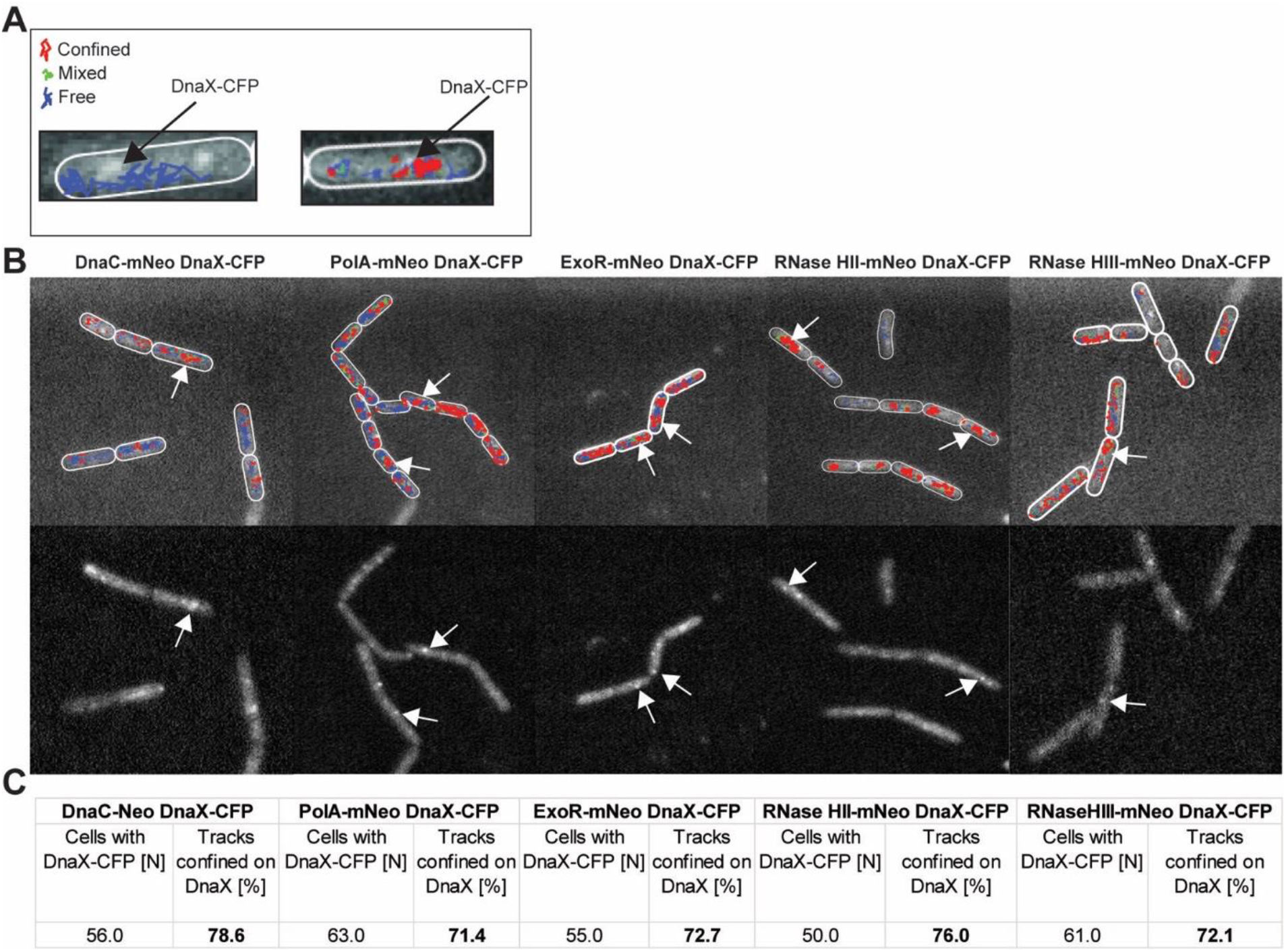
Confinement analyses of replication-associated proteins relative to DnaX-CFP foci. A) example of two cells, one in which freely diffusing tracks are shown in blue, and one where confined tracks overlapping with a DnaX-CFP focus are shown. B) upper panels show an overlay of confinement analyses of DnaC-mNeo, PolA-mNeo, ExoR-mNeo, RNase HII-mNeo, and RNase HIII-mNeo, in relation to the foci of replication forks (DnaX-CFP, lower part). More than 50 cells per strain were analysed. C) Table of percentage counts. The cells with visible DnaX-CFP signal are indicated (N). Cells with confined tracks at the replication fork are indicated in percent [%]. The tracks are divided into confined (red), in transition (green) and free (blue).

These data strongly suggest that Okazaki fragment maturation involves the recruitment of at least four enzymes to the replication forks in *B. subtilis*. Because we wished to obtain additional evidence for this idea, we tested if enzymes would be more strongly engaged as replication forks become stalled, e.g. in the response to UV irradiation, which of course also induced DNA repair events at many other sites on the chromosome.

### PolA and RNase HIII show a change in mobility in response to UV light-induced DNA damage

To test the influence of DNA damage on the dynamics of RNases HII and HIII and to identify a possible involvement in the UV repair system, the dynamics of the proteins under UV-damage of DNA was determined. Therefore we induced DNA damage *via* crosslinks using UV light (Duigou *et al*., 2005), using a treatment of 60 J m-2. Please note that for this analysis, diffusion constants for each protein were determined by a simultaneous fit, in order to better compare changes in population size. Interestingly, only the population sizes of PolA-mNeo change significantly. After UV treatment, there was an increase in the static population from 30.2% to 43.2%, with a concomitant decrease of the high-mobile fraction (Fig 5A and B). ExoR-mNeo, RNase HII-mNeo and RNase HIII-mNeo did not show any significant changes in their dynamics in response to UV treatment. This is somewhat contradictory to earlier reports of our group, in which significant changes were also detected for ExoR (Hernández-Tamayo *et al*., 2019). We will come to this point later.

**Figure 5:**
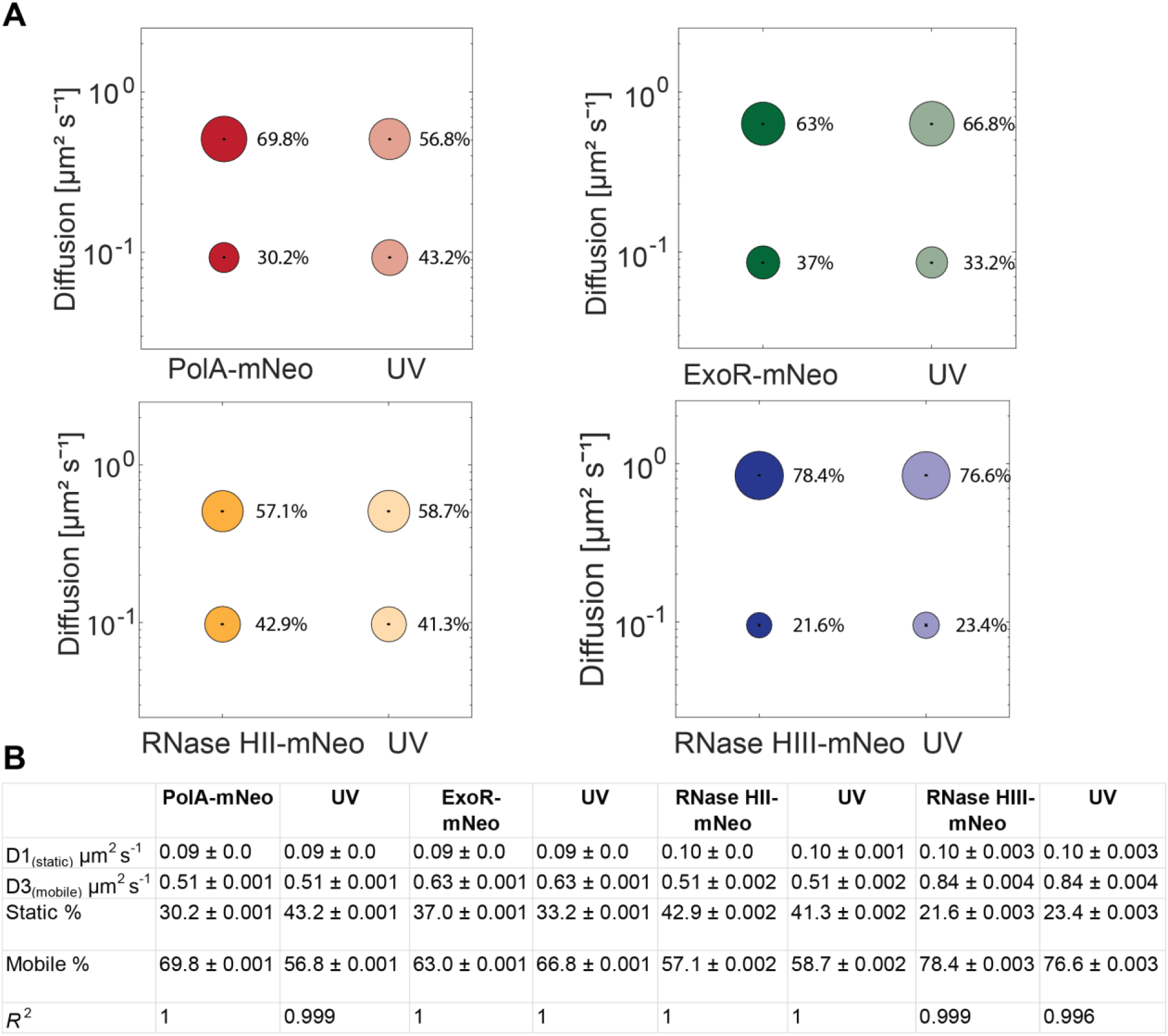
Analyses of protein dynamics via Single molecule tracking with or without UV-treatment. (A) Bubble blots show diffusion constants of replication Proteins and fractions sizes for static and mobile molecules with and without treatment with UV-light. (C) Diffusion constants and percentages of static and mobile molecule fractions. Values were fitted using non-linear least-square fitting, R2.

We employed proximity determination as a second means to detect changes in protein dynamics/localization after induction of DNA damage. Following UV treatment, PolA-mNeo showed a shift of the probability of tracks towards the “0” position (Fig 3C), indicating even stronger association with DnaX and thus replication forks. For ExoR-mNeo of RNase HII-mNeo, there was no noticeable change of distance to DnaX-CFP foci with and without UV light treatment (Fig 3D and E). However a considerable change was seen for RNase HIII-mNeo. Without treatment, the distribution showed a peak at about 0.4 μm, and a maximum probability of occurrence of 0.05. After UV treatment, there was an almost threefold increase in abundance at a distance of 0.4 μm from 0.05 to about 0.14 (Fig 3F). This finding argues for a higher accumulation of RNase HIII-mNeo at the replication forks during DNA damage by UV. For reference, DnaC-mNeo showed a peak at about 0.4 μm with a probability frequency of 0.075 (Fig. 3B).

### Inhibiting PolC activity leads to a strong effect on the localization of RNase HII and HIII

Because there were no detectable changes for single molecule dynamics of RNase HII-mNeo or RNase HIII-mNeo during UV stress, we employed a third treatment, to arrest DNA replication via inhibition of DNA polymerase PolC, using 6(p-hydroxyphenylazo)-uracil (HPUra), which reversibly binds to and inhibits DNA polymerase PolC, thereby completely blocking progression of replication (Brown, 1970), as opposed to slowing down replication due to the necessity to repair based dimers in response to UV irradiation. In this case, we used mVenus fusion strains, for which we did not observe any differences with respect to single molecule dynamics. Fig. 5 shows corresponding heat maps for RNase HII-mV and RNase HIII-mV, which display a visually clear difference in the preferred localization of RNase HII-mV in the cell (Fig 6 A): during exponential growth, a high concentration of tracks is found in the central area of nucleoids, where replication takes place, which is lost during replication arrest. With regard to population size, RNase HII-mV shows a considerable decrease in the static population from 49.5% to 40.2% in response to HP Ura treatment, indicating that a large pool of substrate-binding sites is abolished. Interestingly, although RNase HIII did not show a strong visual alteration in the heat maps of tracks (Fig. 6A), the RNase showed a pronounced decrease in the static population, from 53% to 29.7%. The mobile populations increased accordingly (Fig 6 B, C).

**Figure 6:**
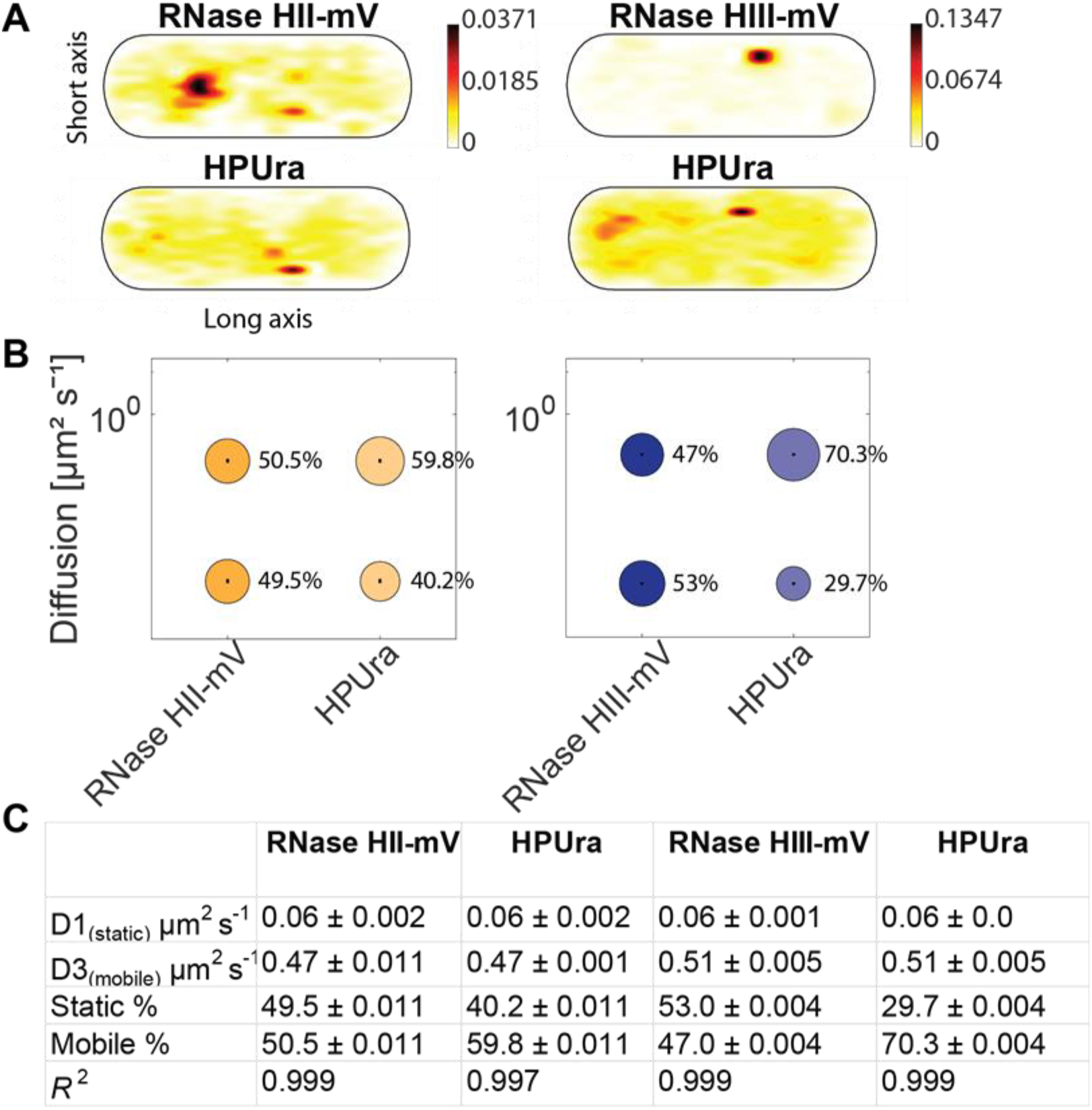
Analyses of protein dynamics under treatment of HPUra via single molecule tracking. (A) Heat maps of single-molecule localization of RNase HII and HIII with and without HPUra (15mg/ml) in a medium-size *Bacillus subtilis* cell. The distribution of tracks is indicated by a color shift from yellow (low probability) to black (highest probability). (B) Bubble blots show diffusion constants of RNase HII/HIII and fractions sizes for static and mobile molecules under the treatment of HPUra. (C) Diffusion constants and percentages of static and mobile molecule fractions. Values were fitted using non-linear least-square fitting, R2 values for each condition are stated.

Thus, a block in replication fork progression alters dynamics of both type H RNases, strongly supporting the idea that they are intimately involved in the processing of Okazaki fragments.

### Lack of ExoR affects RNase HIII dynamics

Previous work has shown that double mutation of *rnhC* and *exoR* results in a phenotype that is lethal at 25°C (Randall *et al*., 2019). We investigated a possible influence of the proteins on each other’s activity by analyzing the dynamics of each in strains carrying the respective deletion of the other gene (ExoR-mVenus *ΔrnhC*, RNase HIII-mV *ΔexoR*). In case both proteins are recruited to replication forks via specific protein/protein interaction with proteins present at the forks at all times, we would not expect strong alterations in e.g. static populations representing DNA-bound molecules, while independent recruitment due to substrate availability might lead to detectable changes in DNA-bound, slow mobile/static, states.

For these analyses, we projected all tracks showing confined motion from the three biological replicates into an average size cell of 3 × 1 μm size (“confinement heat map”, *B. subtilis* cells are on average 0.75 μm wide and 2 to 4 μm long). To achieve this, tracks were sorted into those that stay within a radius of 120 nm, determined as three times our localization error, for at least 6 consecutive steps (confined motion), and into those that show large displacements, indicative of free diffusion (Hinrichs *et al*., 2022).

For ExoR-mV, as well as RNase HIII-mV with and without deletions, there were inconsiderable difference in the pattern of confined tracks at 30°C (Fig. 7A). Incubation at 37°C, where DNA replication occurs at maximum speed, led to an increase in the degree of confined motion relative to 30°C (Fig 7A). While deletions of *rnhC* or *exoR* did not result in clear changes in the heat maps, considerable changes were observed in the dynamics of the proteins. For ExoR-mV, the static fraction decreased from 47.7% to 36% in the deletion background of *rnhC*, compared to wild type cells (Fig. 6B). Conversely, RNase HIII-mV *ΔexoR* cells showed an increase in the static population during incubation at 37 degrees (43.3% to 51.3%) (Fig. 7B and C). Western blotting was performed to determine the expression levels of the proteins (Fig S2). No significant changes were found, showing that changes in population sizes are due to changed binding behavior (i.e. engagement in slow diffusion) of existing molecules.

**Figure 7:**
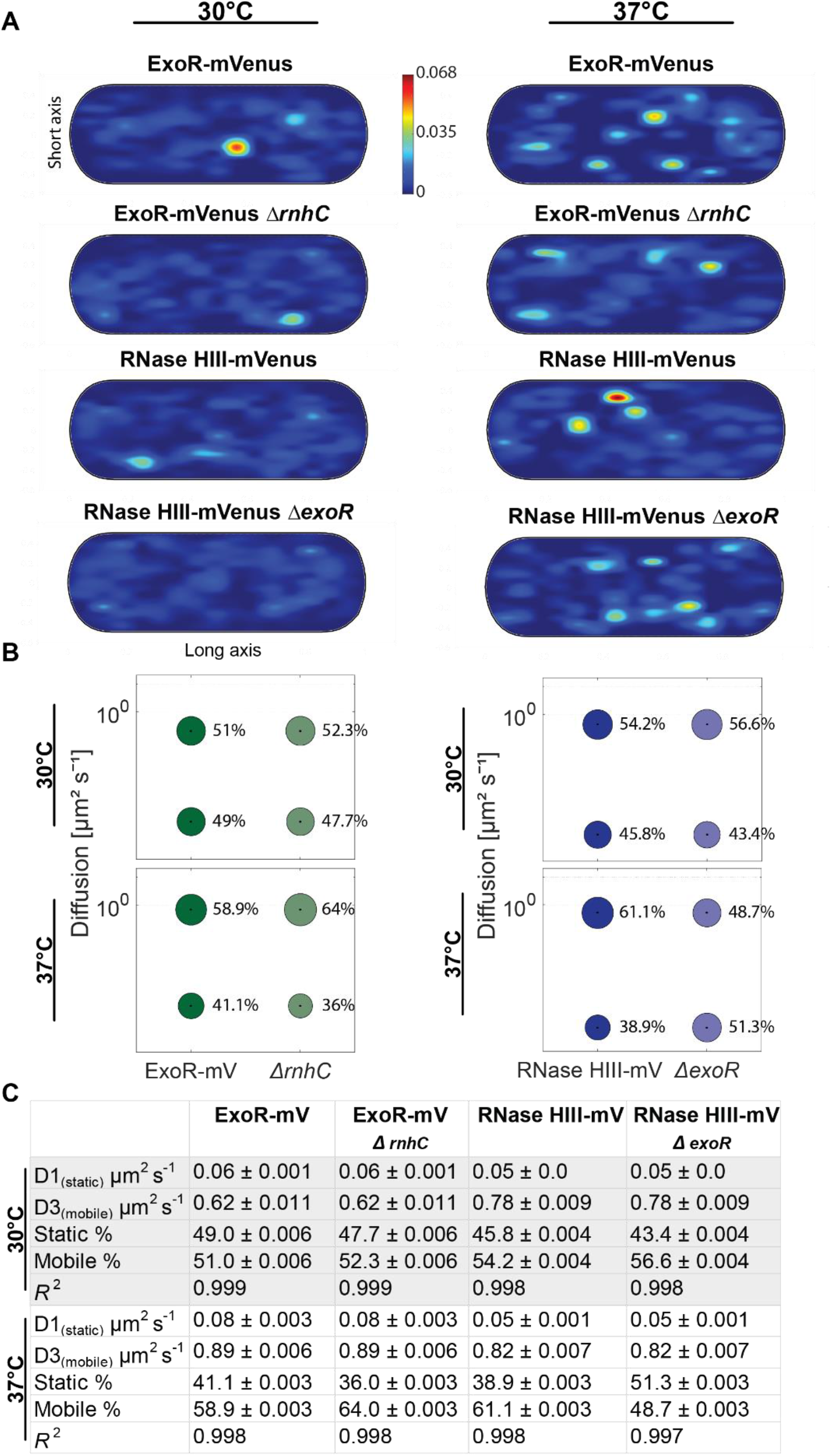
Analyses of protein dynamics via Single molecule tracking at 30°C and 37°C. (A) Confinement maps of potential replication proteins by 30°C and 37°C. Plots of heat map of confined tracks, projected into a standardized *B. subtilis cell*. (B) Bubble blots show diffusion constants of replication Proteins and fractions sizes for static and mobile molecules at different temperatures (30°C and 37°C). (C) Diffusion constants and percentages of static and mobile molecule fractions. Values were fitted using non-linear least-square fitting, R2 values for each condition are stated.

While we can not show where changes in slow mobility occur in a decisive manner, we favour the view that for RNase HIII, lack of ExoR provides more substrate sites, suggesting that RNase HIII is recruited based on the occurrence of DNA/RNA hybrids rather than via specific protein interactions. For ExoR, changes are more difficult to interpret. A reduction of slow mobile/static molecules from 41 to 36% represents a change of 12.2%, which is close to what we would regard as noise in biological systems, and may thus rather be a “no-change” phenotype.

## Discussion

Incorporation of short stretches of RNA as primers for lagging strand synthesis requires the activity of enzymes that remove RNA from DNA/RNA hybrids in all organisms. In eukaryotic cells, RNase H2 and Exo1 together with the flap endonucleases Fen1 and Dna2 have been reported to be involved in Okazaki fragment maturation (Liu *et al*., 2017), while in *E. coli*, it is believed that DNA Pol 1 and RNase HI take over this task. In the Gram positive model organism *B. subtilis*, RNase HIII, exonuclease ExoR and DNA polymerase PolA have been proposed to be involved in replication, based on biochemical and genetic data (Randall *et al*., 2019). It has also been suggested that RNase HIII acts as an important supporter of PolA in the maturation of Okazaki fragments (Randall *et al*., 2019): it has been shown that RNase HIII influences endonucleolytic cleavage of RNA in RNA-DNA hybrid molecules in the processing of R-loops and maturation of Okazaki fragments. Additionally, the absence of both H-type RNases, HII and HIII leads to a synthetic slow-growth phenotype (Yao *et al*., 2013), indicating that also RNase HII may be involved. However, the exact *in vivo* function of RNase HII is not known except for the possible removal of single rNMPs incorporated into DNA by DNA polymerases (DNA replication (Yao *et al*., 2013, Randall *et al*., 2018). We sought to provide *in vivo* evidence to answer the question, which enzymes are truly involved in RNA removal from replication forks. Using single molecule tracking, we could show that RNase HII, RNase HIII, as well as ExoR and PolA arrest at replication forks with a frequency that is close to that of molecules of replicative helicase DnaC. In response to DNA damage induced via UV stress, we found increased activity of RNase HIII and Pol A at the forks.

Single molecule trajectories for all four enzymes could be explained by assuming two populations having distinct diffusion constants. Based on the idea that DNA polymerases or RNases detecting DNA/RNA hybrids would be in a bound state, where there is little diffusion (basically that displayed by DNA strands), the slow mobile population should represent enzymatically active enzymes bound to DNA, while the high-mobile fraction should represent diffusing molecules. Diffusion constants for all proteins were quite low in comparison to freely diffusing enzymes (Rotter *et al*., 2021), and PolA, ExoR and RNase HII showed clear nucleoid staining in epifluorescence microscopy. Heat maps of all single molecule trajectories showed a clear enrichment of tracks at central places, even within nucleoid areas, similar to those of DNA polymerase C, and very different from the more peripheral pattern of RNase J1, which is involved in RNA degradation within the membrane-associated RNA degradosome. These data show that like DNA transcription factors (Stracy *et al*., 2021), or sequence-specific DNA methylases (Negri *et al*., 2021), H-type RNases and ExoR mostly employ constrained motion through the nucleoids, diffusing between DNA strands to find binding targets, which would explain the rather low mobility of the high-mobile fraction.

RNase HII and HIII belong to the RNase H enzyme family, which are responsible for the identification and cleavage of RNA-DNA hybrids (Cerritelli & Crouch, 2009, Ohtani *et al*., 1999b). The formation of R-loops and the resulting DNA-RNA hybrids is a known stress response. This stress can be caused by cell wall damage, osmotic stress, oxidative damage, but also DNA damage, and can lead to genomic instability and replication arrest (Gan *et al*., 2011, Lin & Pasero, 2012). Using confinement analysis relative to the replication forks, we were able to clearly show that all four analysed enzymes have a very similar percentage of confined tracks in the direct proximity of the replication fork, similar to DNA helicase (DnaC). Confined tracks show dwell events for at least 100 ms, likely reflecting enzymatic activity in the DNA-bound form. Based on distance determination between molecule trajectories and replication forks, we could clearly see an abundance of RNase HIII at the replication forks. Interestingly, during the influence of DNA damage (UV), RNase HIII became more enriched at sites close to the forks, possibly reflecting its published involvement in R-loop processing in *Bacillus subtilis* (Lang *et al*., 2017). Interestingly, RNase HII also showed a high degree of spatial proximity to sites of DNA replication, strongly suggesting it is likewise involved in Okazaki fragment maturation (based on the notion that incorporation if RNA nucleotides by DNA polymerases is a very rare event). In contrast to RNase HIII, HII did not show any stronger shift to the forks after inducing DNA damage by UV. PolA showed a stronger engagement with forks following UV irradiation, as was reported before (Hernández-Tamayo *et al*., 2019). In contrast to this study, we found PolA even strongly associated with DNA replication sites even before induction of DNA damage, and similarly for ExoR. We believe that these differences are due to larger sample sizes in this work, showing significant localization even during normal growth. Likewise, the mNeo fluorophore used can lead to lower artifacts, as it has a higher lifetime, higher brightness, with a shorter maturation time in contrast to mVenus (Shaner *et al*., 2013). The improvement of the fluorophore in combination with the higher sample size can lead to deviations.

In order to use a third means to show involvement of RNase HII and HIII, we tracked the fusion proteins during a replication block by HPUra, where the activity of PolC is inhibited. We observed strong differences in mobilities and localization pattern. Both RNases became more dynamic, with RNase HIII showing the largest change from static to mobile diffusion. For RNase HII, we observed a strong relocalization of molecules from central places on the nucleoids towards more peripheral sites. Overall, our *in vivo* analysis provide evidence that both proteins are actively involved at replication forks, With RNase HIII playing a more pronounced role than RNase HII. Similarly, only RNase HIII together with DNA polymerase I (PolA) seems to show significant responses to UV stress, underlining the already assumed interdependence of the two proteins. It could be shown that RNase HIII supports PolA in the maturation of Okazaki fragments (Randall *et al*., 2019). Interestingly, RNase HII as well as ExoR show very similar dynamics and localizations in the cell. RNase HII is known to remove single ribonucleoside monophosphates (rNMPs) during DNA replication (Yao *et al*., 2013) The function is important for the cell because DNA is more stable than RNA. Resulting rNMP residues in DNA could lead to spontaneous strand breaks. ExoR and endoribunclease RNase HII showed very similar dynamics and localization, but it is unclear if they may have similar functions, in line of their synthetic lethality with PolA or RNase HIII, respectively (Thomaides *et al*., 2007). However, RNase HII and HIII treated with HPUra demonstrated a response to replication blockage. So, the single molecule tracking of these treated strains has revealed although that RNase HII and RNase HIII appear to have distinct functions *in vivo*, the analysis suggests that RNase HII may have a supporting effect in the maturation of Okazaki fragments and might be a RNase involved in this process.

We addressed the question whether enzymes may be specifically recruited to forks in order to remove RNA primers, or whether these sites are found by a diffusion/capture mode. We therefore analysed the dynamics of ExoR and RNase HIII in the corresponding deletion backgrounds, as a cold-sensitive phenotype for double mutations of *exoR* and *rnhC* were observed (Randall *et al*., 2019). The dynamics of the proteins did not show notable changes at 30°C, but at 37°C, RNaseH III-mV revealed a remarkable increase in the static population in the absence of *exoR*. ExoR-mV showed a slight decrease in the static population in the deletion background of *rnhC*. Clearly the lack of ExoR leads to mode binding events of RNase HIII, based on its higher abundance in the low mobility state, apparently leading to a takeover of RNA nucleotide removal by RNase HIII. These data suggest that RNAse HIII shows a diffusion/capture mode of recruitment at replication forks, besides possible, unknown direct protein/protein interactions.

**Figure 7:**
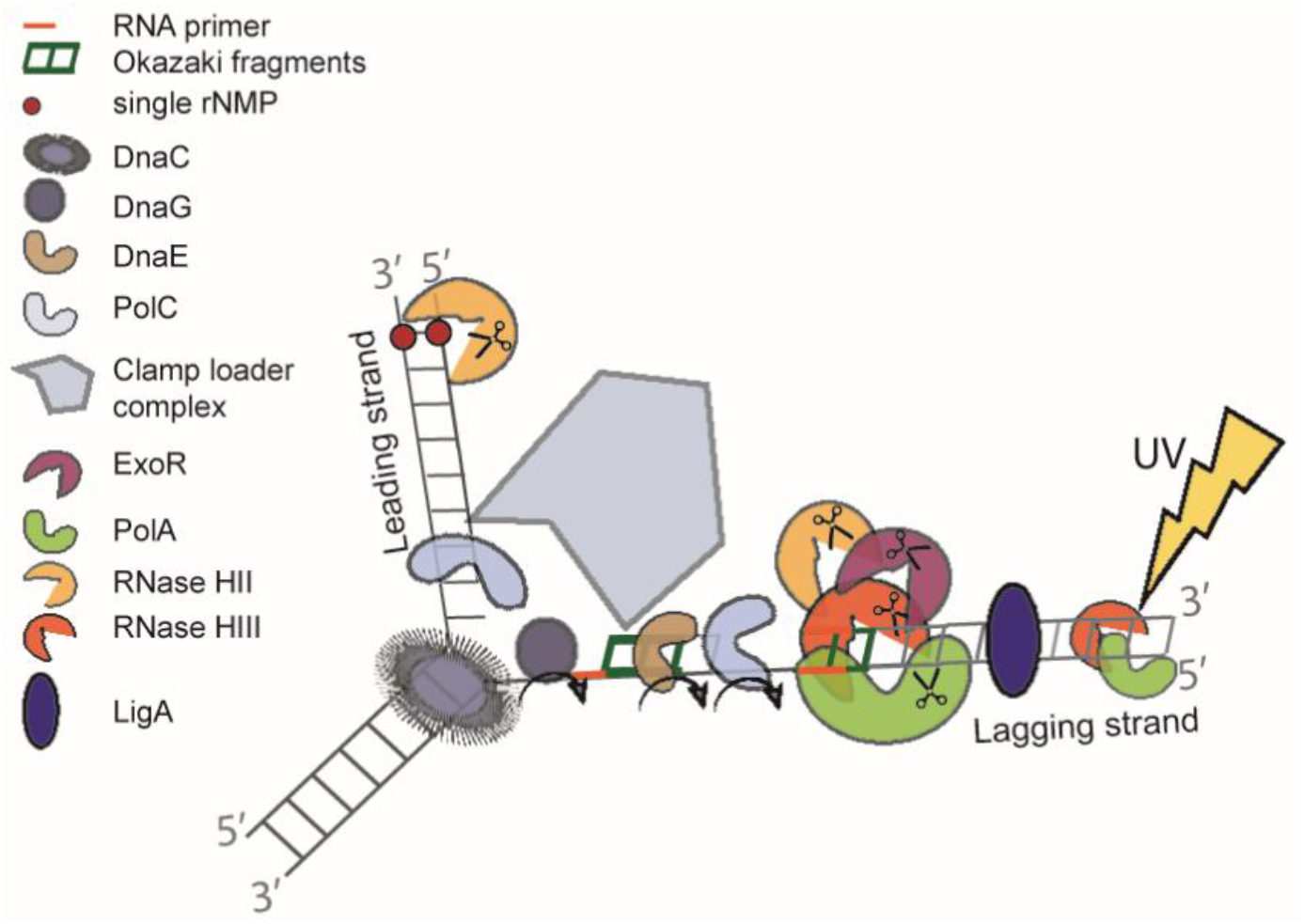
Graphical illustration of replication in *Bacillus subtilis* during UV stress. At the replication fork, a DNA helicase (DnaC) precedes the DNA synthesis machinery and unwinds the exist duplex parental DNA in cooperation with the single stranded binding proteins (SSB). On the leading strand (5 to 3 direction), replication proceeds continuously via the replisome. In contrast, on the lagging strand, DNA replication is performed discontinuously by synthesizing and assembling short Okazaki fragments. DNA primase (DnaG) is required for the formation of RNA primers. During DNA template replication, an RNA primer is removed either by the 5-bis-3 exonuclease activity of ExoR and Okazaki fragments are matured via PolA, RNase HII and RNase HIII. Afterwards DNA ligase LigA fuses matured Okazaki fragments.

With the help of previous work (Randall *et al*., 2019, Yao *et al*., 2013, Randall *et al*., 2018, Patlán *et al*., 2019, Jameson & Wilkinson, 2017) and our *in vivo* analyses, we conclude that *B. subtilis* has replication forks that strongly deviate from textbook knowledge on bacterial replication forks (Fig 7). Initiation of replication leads to loading and ensuing unwinding activity of the double-stranded DNA by the DNA helicase DnaC. The leading strand (5 to 3 direction) is then continuously processed by DNA polymerase C holoenzyme. For the replication of the lagging strand (3 to 5 direction), discontinuous processing occurs. Small RNA primers start each Okazaki fragment, which is synthesized by the primase (DnaG). DNA polymerase E, extends RNA primers by some DNA bases, and hands over synthesis of the lagging strand to DNA PolC. DNA ligase (LigA) ligates all fragments of the lagging strand at the 3’ end. At least four specific exo-or endonucleases are used to remove RNA primers from the lagging strand. A known one is DNA polymerase I (PolA), which removes these primers with its exonuclease function (5’-3’ exonuclease activity) together with the endoribonuclease RNase HIII (Randall *et al*., 2019). In a likely redundant manner, RNaseH II and ExoR also remove RNA primers, while only RNase HIII and PolA show increased recruitment are suitable for UV damage and could be a part of the SOS response in *B. subtilis* (Lenhart *et al*., 2012). However, it can be assumed that RNase HIII has functions outside of replication forks, because it shows areas of constrained motion at many sites on the nucleoids (Fig. 6A).

Altogether, our work clarifies functions for four enzymes removing DNA/RNA hybrids from replication forks, which may be a unique multitude of proteins for *B. subtilis*. It will be interesting to investigate if some or many other bacterial species also distribute RNA removal during replication onto many enzyme’s shoulders.

## Acknowledgments

This work has been supported by the LOEWE funded consortium MOSLA (state of Hessen).

## Author contribution

RH has performed all experiments, and co-wrote the manuscript. PLG supervised experiments, conceived of the study, and co—wrote the manuscript.

## Data availability statement

All data are within the manuscript. Raw data for tracking will be made available after reasonable request at the corresponding author

## Supplementary Material

**Figure S1:**
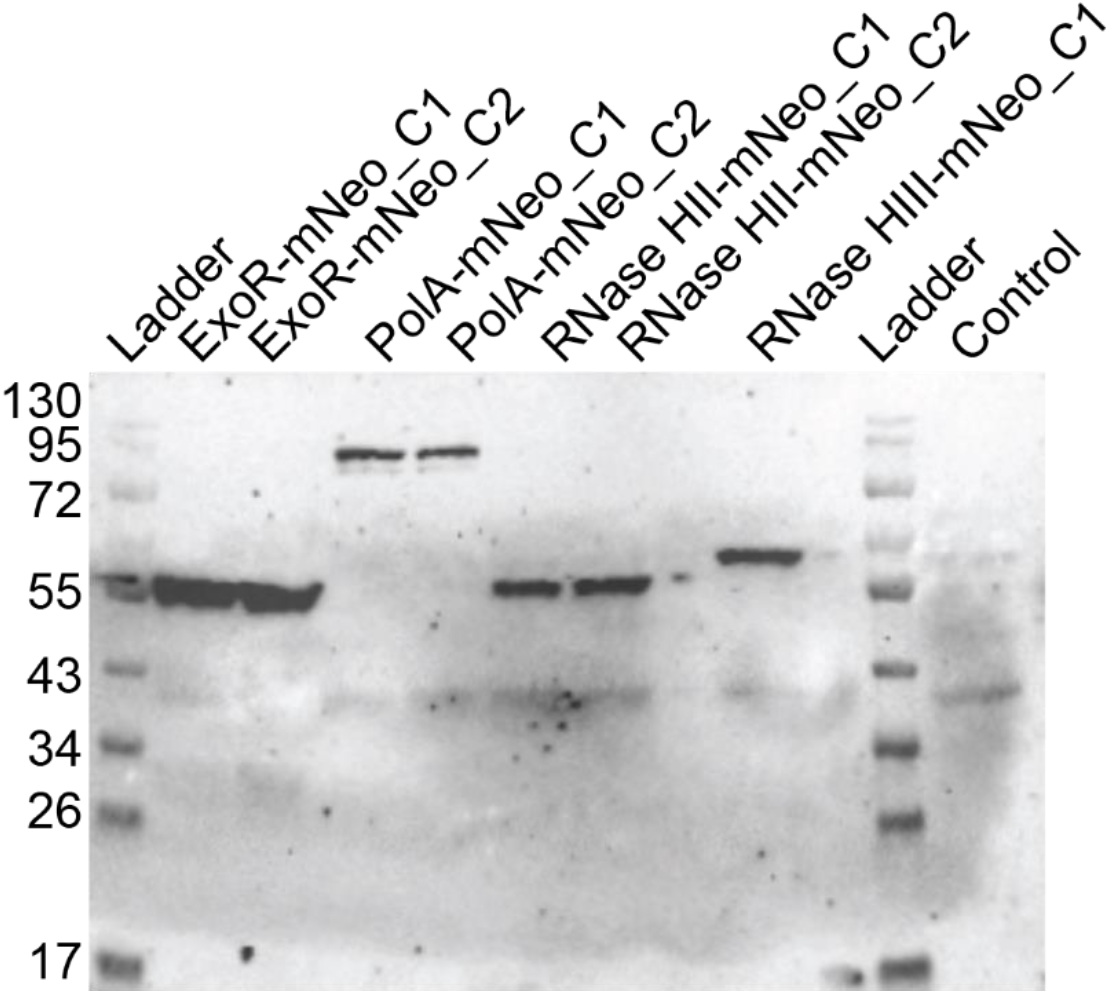
Western blot of potential replication proteins. Western blots showing mNeo fusion expressed from native locus. Total cell extracts from exponentially growing cultures (LB) were used. Two different clones are shown in each case (C1/C2). The ExoR-mNeo fusion (59.7 kDa), PolA-mNeo (125.8 kDa), RNaseHII-mNeo (55.1 kDa) and RNaseHIII-mNeo (60.8 kDa) contains the mNeongreen polypeptide (26.9 kDa). All strains were detected via mNeongreen-antiserum. As a control strain, the *Bacillus subtilis* BG214 was used.

**Figure S2:**
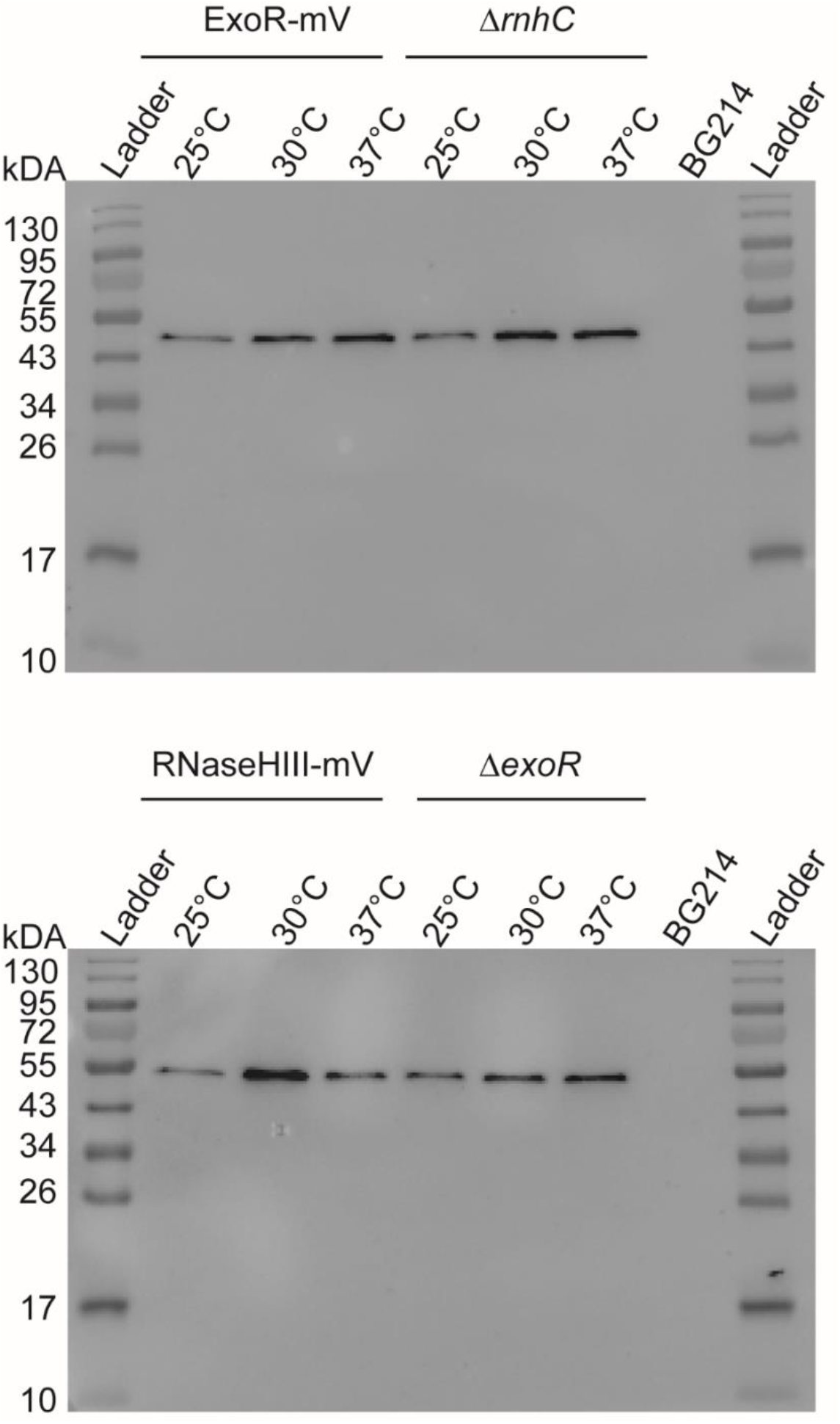
Comparison of expression level under deletion conditions and different temperatures. Western blots showing mVenus fusion expressed from native locus. Total cell extracts from exponentially growing cultures (LB), incubated at different temperatures (25°C, 30°C, 37°C) were used. The ExoR-mV fusion as well as the deletion of rnhC (59.7 kDa), RNaseHIII-mV (60.8 kDa) and the delition of exoR contains the mVenus polypeptide (26.9 kDa). All strains were detected via mVenus-antiserum. As a control strain, the Bacillus subtilis BG214 was used.

